# *RPS28B* mRNA acts as a scaffold promoting *cis*-translational interaction of proteins driving P-body assembly

**DOI:** 10.1101/696823

**Authors:** Nikita Fernandes, J. Ross Buchan

## Abstract

P-bodies (PBs) are cytoplasmic mRNA-protein (mRNP) granules conserved throughout eukaryotes which are implicated in the repression, storage and degradation of mRNAs. PB assembly is driven in part by proteins with self-interacting and low-complexity protein domains. Non-translating mRNA is also required for PB assembly, however no studies to date have explored whether particular mRNA transcripts are more critical than others in facilitating PB assembly. A previous genome-wide microscopy screen in yeast revealed that *rps28bΔ* (Ribosomal protein subunit-28B) mutants do not form PBs under normal growth conditions. Here, we demonstrate that the *RPS28B* 3’UTR is important for PB assembly, consistent with the fact that this is a known binding site for the PB assembly protein Edc3. However, expression of the *RPS28B* 3’UTR in isolation is insufficient to drive normal PB assembly. Intriguingly, chimeric mRNA studies revealed that Rps28 protein, translated *in cis* from an mRNA bearing the *RPS28B* 3’UTR, physically interacts more strongly with Edc3 than Rps28 protein synthesized *in trans*. This Edc3-Rps28 interaction in turn also facilitates PB assembly. In summary, our work indicates that PB assembly may be preferentially nucleated by specific RNA “scaffolds”, which may be a common theme in RNP granule assembly. Furthermore, this is the first description in yeast to our knowledge of a *cis*-translated protein interacting with another protein in the 3’UTR of the mRNA which encoded it, which in turn has functional consequences for assembly of cellular structures.

## Introduction

Post-transcriptional processes are crucial regulators of gene expression. In the cytoplasm, mRNAs can have multiple fates. They can either be actively translated or be subject to translational repression followed by storage or decay. All of these non-translating mRNP states have been associated with P-bodies (PBs)[1,2].

PBs are membrane-less cytoplasmic RNA granules conserved from yeast to humans[3][4]. They are present in all cells growing under normal growth conditions but typically increase in size and number during stress[5]. They are composed of non-translating mRNAs and harbor numerous mRNA decay proteins including many involved in 5’-3’ mRNA decay, as well as nonsense-mediated decay (NMD) proteins[6]. miRNA and siRNA-associated proteins also localize in PBs[7]. While earlier understanding of PB composition largely derived from fluorescence microscopy approaches, recently, PBs were isolated from HEK293 cells and subject to mass spectrometry and RNA-Seq analysis, greatly increasing our understanding of the compositional makeup of these structures[8].

Systematic experiments posit that PB assembly is a two-step process requiring the generation of a pool of non-translating mRNP complexes that then interact via proteins that harbor self-interaction and low complexity domains (LCDs). First, supporting the role of non-translating mRNPs in PB assembly, treatment of cells with cycloheximide, which traps mRNAs in polysomes, inhibits PB assembly[5]. Conversely, inhibition of translation initiation[5] or treatment with puromycin[9], all increase the amount of non-translating mRNPs, and result in an increase in PB assembly. Furthermore, *in vitro* RNase treatment of semi-purified PBs result in their disassembly[5]. Second, supporting the role of specific protein interactions in PB assembly, deletion of genes including Edc3, Pat1 and Lsm4 lead to defects in PB assembly [10][11]. Edc3 has a self-interaction domain (Yjef-N) and Lsm4 has a glutamine/asparagine (Q/N) rich low complexity domain that are both implicated in PB assembly[10]. Furthermore, Edc3 directly interacts with multiple PB proteins like Dcp2 and Dhh1, whereas Pat1 can additionally bind to Xrn1, Dcp2, the Lsm1-7 and Ccr4-Not complexes[12,13][14]. By virtue of these multivalent interactions with numerous PB proteins, Edc3 and Pat1 act as protein scaffolds in PB assembly. Thus, like other mRNP granules, PB assembly is driven by multiple proteins, with no single protein seemingly essential for PB formation[15].

To date, knowledge about PB assembly has largely been gleaned from candidate gene analyses. However, an unbiased genetic screen coupled to live cell yeast microscopy to identify genes that alter PB assembly identified *rps28bΔ* as having a severe defect in PB formation under normal growth conditions[16]. Rps28 is a protein of the 40S ribosomal subunit and binds near the mRNA exit tunnel. In yeast it is encoded by paralogous genes *RPS28A* and *RPS28B*. Rps28a and Rps28b are identical except for an S3N change. However, they differ significantly with respect to their mRNA 3’UTRs. *RPS28B* mRNA possesses an unusually long 3’ UTR of ~643nts. The 3’UTR also harbors a stem loop structure, thought to bind Edc3[17], that has been extensively studied for enabling an auto-regulatory circuit that regulates *RPS28B* mRNA and protein levels [18]. Specifically, it is proposed that high Rps28 protein levels, generated from either *RPS28A* or *RPS28B* mRNA, leads to Rps28-Edc3 binding. This is thought to promote recruitment of decapping proteins to the mRNA, and drive deadenylation-independent decapping and decay of *RPS28B* mRNA[17–19]. Thus, Rps28 protein levels are thought to regulate *RPSB28B* mRNA abundance, thus helping maintain a homeostatic balance of Rps28 protein. Interestingly, in normal growth conditions at mid-log, Rps28a protein is reportedly 11 fold more abundant than Rps28b, but *RPS28A* mRNA levels are only 50% greater than that of *RPS28B* [18,20], suggesting that *RPS28B* mRNA is less translationally active, for reasons that remain unknown.

While deletion of *RPS28B* could cause a decrease in Rps28 protein levels resulting in PB assembly defects, we hypothesized that the mRNA itself might also have a role as a novel mRNA scaffold driving PB assembly for two reasons. 1) *RPS28B* mRNA 3’UTR interacts with Edc3, a major PB assembly factor[17,18]. 2) The *RPS28B* 3’UTR is unusually long. Recent studies have shown that mRNA lengths correlate with their enrichment in RNA granules[21,22]. Furthermore, RNAs themselves can drive liquid-liquid phase separation (LLPS)[23,24]. LLPS is a process thought to facilitate granule formation wherein biomolecules with high valency phase separate once critical local concentrations are attained. While RNAs can accelerate this process *in vitro*[23] (though not always [25]), importantly, no specific mRNA has been identified to uniquely and potently facilitate PB (or stress granule; SG) assembly *in vivo*.

In this work, our data suggests that the *RPS28B* mRNA 3’UTR acts as a novel PB nucleating mRNA scaffold. However, Rps28 protein, translated *in cis* from the *RPS28B* mRNA, is also important for PB assembly. Strikingly, the *cis* translation of Rps28 from the *RPS28B* mRNA with its native 3’UTR is necessary for efficient Rps28-Edc3 protein interaction, which in turn is necessary for PB assembly under normal growth conditions. This work suggests that mRNA scaffolds might be a common theme in RNA granule assembly. More broadly, and consistent with recent work [26][27,28], an under-appreciated role of mRNA 3’UTRs may be enhancement of protein-protein interactions involving nascently encoded proteins and previously 3’UTR-bound binding partners.

## Materials and methods

### Yeast strains and growth conditions

The strains used in this study are described in Supplemental Table 1. The strains knocked out for specific genes were obtained from the Yeast Knockout Collection. Strains were grown on YPD or synthetic media (VWR Glucose 2%, Difco Yeast Nitrogen Base 0.17%, Fisher Ammonium sulphate 5 g/L, appropriate amino acids and nucleotides). All strains were grown at 30°C in shaking water baths. A standard lithium acetate technique was used for yeast transformations. Glucose deprivation stress was applied for 10 min as previously described[11].

### Plasmids

The plasmids used in this study are described in Supplementary Table 1. To construct plasmid pRB224, GFP was amplified from plasmid pRB001 using oligos oRB446 and oRB447. This PCR product along with XhoI digested pRB011 was recombined in yeast via homologous recombination. To make plasmids pRB378, pRB229 and pRB230 (RPS28B 3’UTR truncation constructs1, 2 and 3), oligos oRB396 and oRB445, oRB398 and oRB399, oRB400 and oRB401 were used to carry out linear amplification of the plasmids with deletion of the respective regions. This was followed by PNK treatment and ligation to circularize the plasmids. To make plasmids pRB379, pRB380, pRB381 and pRB382, plasmids pRB225, pRB227, pRB228 and pRB226 were digested with NotI and SalI. Required gel extracted fragments were then ligated into NotI/SalI cut pRS315.

### Microscopy

The screen that identified *rps28bΔ* as having a severe defect in PB formation has been previously described[16]. Analyses of PB assembly was conducted as previously described[29], using a Deltavision Elite (100x objective) followed by image analysis using Fiji software[30]. For every strain, a minimum of 100 cells/per replicate were quantified, with a minimum of 3 biological replicates examined for each microscopy dataset.

### Polysome analysis

Polysome analyses were conducted as previously described[31] with the following minor differences. Cells were either harvested in midlog (OD_600_ 0.3-0.6) or after being subject to 15 min glucose deprivation stress.

### Western Blots

Western blotting was carried out by standard protocols. Protein extracts were loaded onto SDS-polyacrylamide gels. Rps28 was detected by using Rabbit anti-Rps28 (Thermo Fisher Scientific) at a 1:2500 dilution. GAPDH was detected by using Mouse anti-GAPDH (Thermo Scientific) at a 1:25000 dilution. Detection was carried out by using Li-Cor fluorescently labelled secondary antibodies and the Li-Cor Odyssey Imaging system. Secondary antibodies used were IRDye 800CW Goat anti-Rabbit IgG (H + L) and IRDye 800CW Goat anti-Mouse IgG (H + L) to detect anti-Rps28 and anti-GAPDH antibodies respectively.

### Northern Blots

5 ml yeast cultures were grown to midlog (OD_600_ ~0.3-0.6) after which RNA was extracted using a hot phenol extraction protocol described previously[32]. Northern blots were carried out using previously established protocols[33]. 30μg of total RNA was subject to agarose gel electrophoresis (1.25% agarose). mRNAs were visualized by PhosphorImager analysis (Typhoon) of nylon membranes that were probed with 5’ end [^32^P]-radiolabeled DNA probes. *RPS28B* was also specifically visualized with a probe complementary to the *RPS28B* 3’UTR (See supplemental table 1). Image analysis was carried out using Fiji[30].

### qRT-PCR

2 ml yeast cultures were grown to midlog (OD_600_ ~0.3-0.6) after which RNA was extracted using the Trizol RNA extraction method. Briefly, cells were centrifuged at 4000rpm for 15 min followed by resuspension in 1 ml of Trizol reagent. Glass beads were added to the microfuge tubes and cells were disrupted using the Fisher Vortex Genie 2 for 5 min at 4°C. 200μl chloroform was added, vortexed for 15 sec followed by a 5 min incubation at RT. Tubes were centrifuged at 13,300rpm for 5 min at 4°C. Aqueous layer was recovered followed by another chloroform extraction. RNA was precipitated by adding 500μl of isopropanol and incubating on ice for 15 min. Tubes were centrifuged at 13,300rpm for 15 min at 4°C. Pellets were washed with 70% ethanol and re-dissolved in 50μl RNase free distilled water. qPCR was carried out using the SuperScript III Platinum SYBR Green One-Step qRT-PCR Kit as per manufacturer’s instructions using oligos mentioned in supplemental table 1. Control experiments were carried out to assess the specificity of the probes used.

### FISH

FISH was carried out using Stellaris FISH probes for *RPS28B* designed using the Stellaris Probe designer. The FISH experiment was carried out using the manufacturers protocol for *S. cerevisae* with the following modifications. Spheroblasting buffer used had components as mentioned in the immunofluorescence protocol above but dissolved in 1 ml Fixation Buffer. Hybridization was carried out at 32°C. Probes are listed in Supplemental Table 1.

### Immunoprecipitation

400ml culture of OD_600_~0.6 was centrifuged at 3000rpm for 2min followed by transfer to a microfuge tubes, centrifugation at 3000rpm for 1min. Pellets were frozen in liquid nitrogen and stored at −80°C until used. Tubes were thawed on ice followed by addition of 300μl lysis buffer (50 mM Tris, pH 7.4, 1 mM EDTA, 150 mM NaCl, 0.5% NP-40) with fungal-specific protease inhibitor (1Ul/100Ul; Sigma) and 1mM PMSF. Acid washed glass beads were added to 500μl and cells were disrupted by using the Fisher Vortex Genie 2 for 3 min at 4°C followed by resting on ice for 2 min. This was repeated twice. Holes were created at the base of microtubes harboring lysed cells and glass beads using a hot syringe needle, and then placed in 15ml tubes. After centrifugation at 2000g for 2 min at 4°C, supernatant was added to equilibrated Chromotek GFP-Trap-MA beads and rotated on a nutator for 1h. This was followed by 4 washes in the abovementioned buffer (excluding NP-40 from this wash buffer). SDS Sample loading buffer was added, samples were heat at 95°C for 10 min followed by western blotting as mentioned above. Rps28 was detected by using Rabbit anti-Rps28 (Thermo Fisher Scientific) at a 1:500 dilution and GFP was detected by using Rabbit anti-GFP (Abcam) at a 1:5000 dilution.

### EMSA assay

Electrophoretic mobility shift assay was carried out as previously described[34], specifically the protocol for simple interactions. The reaction mixture of ^32^P end-labelled RNA and Edc3 was incubated for 1 h at 30°C. Edc3 (N-terminal SNAP and C-terminal His6 tagged) was a gift from the Parker lab.

## Results

### RPS28A and RPS28B are both important for PB assembly

A previous genome-wide microscopy screen to identify genes that affected PB and stress granule assembly in yeast identified *RPS28B* as a gene that, when deleted, conferred a severe defect in PB assembly under normal growth conditions[16]. However, deletion of its paralog *RPS28A* was not tested. We found that in addition to *rps28bΔ, rps28aΔ* strains are also severely defective in PB assembly under normal growth conditions, with *rps28bΔ* showing ~100% and *rps28aΔ* showing ~70% reduction in number of PBs per cell respectively as assessed by Edc3 foci (Figure 1A-B). We also examined PB assembly in the above strains during glucose deprivation stress. As expected, PBs increased in size and number in WT cells under stress as previously observed, and although *rps28aΔ* and *rps28bΔ* strains did not completely block PB formation, a significant inhibitory effect was still observed, similar to that of an *edc3Δ* strain in magnitude[11] (Figure 1C-D). Impaired PB assembly in *rps28aΔ* and *rps28bΔ* strains was also observed when other PB markers like Dhh1 and Dcp2 were used to assess PB formation (Figure S1A-B). The PB assembly defect is also unlikely due to general ribosome impairment caused by absence of any given ribosomal protein, as strong PB assembly defects were not seen in multiple other viable ribosomal protein deletion mutants in the original screen[16], or mutants re-assessed here under normal growth conditions (Figure S1C-D). Edc3-GFP protein levels in WT, *rps28aΔ* and *rps28bΔ* strains also showed no differences, thus arguing impaired PB assembly is not an artefact of altered PB marker expression (Figure S1E). Finally, WT and *rps28aΔ* growth rates are almost equal, with *rps28bΔ* strains showing only a minor growth defect (Figure S1F). In summary, deletion of *RPS28A* and *RPS28B* genes results in severe impairment in PB formation under normal growth and stress conditions.

**Figure 1.**
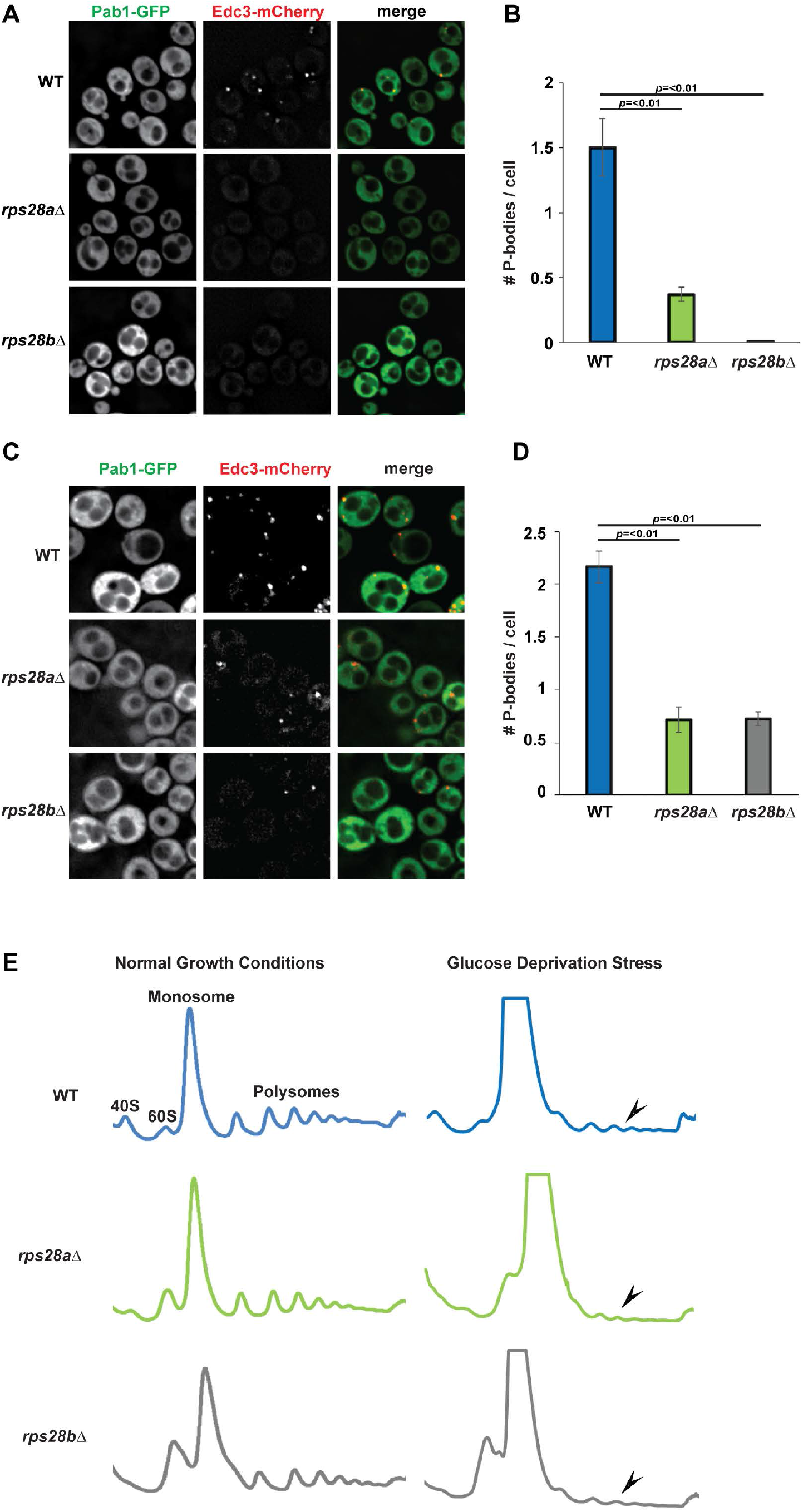
RPS28A and RPS28B are important for PB assembly. A) Log-phase wild-type *S.cerevisiae* BY4741 (WT*), rps28aΔ* and *rps28bΔ* strains were transformed with pRB001 expressing both Pab1-GFP (SG marker) and Edc3-mCh (PB marker) and examined for the presence of PB foci. B) Quantification of A; average number of PBs per cell. Data generated from 3 biological replicates with mean ± s.d shown. An ANOVA with Dunnetts post-hoc test was used to assess significance. C) As in A, except cells were subject to 10minute glucose deprivation stress. D) Quantification of C; average number of PBs per cell. Data generated from 3 biological replicates with mean ± s.d shown. An ANOVA with Dunnetts post-hoc test was used to assess significance. E) *rps28aΔ* and *rps28bΔ* strains are not defective in global translation repression. Log-phase WT*, rps28aΔ* and *rps28bΔ* strains growing under normal growth conditions and under 10 min of glucose deprivation stress were subject to polysome analysis. Data is representative of observations from three biological replicates.

### rps28aΔ and rps28bΔ strains are not defective in global translation repression

Prior literature suggests that PB assembly defects can be due to either loss of assembly factors (e.g. Edc3, Lsm4[10]), or a defect in translation repression or polysome dissociation, which hampers the accumulation of non-translating mRNPs that are necessary for PB assembly[5,35]. To address this second possibility, we subjected *rps28aΔ* and *rps28bΔ* strains to polysome analysis under normal growth and glucose deprivation stress, which elicits a strong translational repression response (Figure 1E); this was previously used to identify Pat1 and Dhh1 as factors affecting translational repression, the first event in PB assembly[35]. In normal growth conditions, polysome abundance in WT, *rps28aΔ* and *rps28bΔ* strains is approximately equal. Following glucose deprivation, WT, *rps28aΔ* and *rps28bΔ* strains all exhibited a strong polysome collapse, indicating no significant defects in global translation repression under stress. Of note, relative to WT, the *rps28aΔ* and *rps28bΔ* strains show a small 40S peak and a large 60S peak, both in normal and stress conditions. This is in keeping with earlier studies that have shown that ribosomal proteins gene deletions lead to aberrancies in 40S and 60S peaks[36,37]. In summary, PB assembly defects in the *rps28aΔ* and *rps28bΔ* strains are unlikely to be due to a general defect in translation or translation repression and suggests a defect at a later step of PB assembly.

### Rps28 protein levels do not account for PB assembly defects in rps28aΔ and rps28bΔ strains

In principle, *rps28aΔ* and *rps28bΔ* strains could affect PB assembly due to decreased levels of Rps28 protein. Indeed, Rps28 interacts with Edc3, Pat1, Lsm2/4/8 and Pby1 PB proteins as assessed by yeast two hybrid assays[38,39]. However, Rps28b protein reportedly only accounts for ~8% of the Rps28 protein population[20,40,41], and yet *rps28bΔ* strains exhibit a striking PB assembly defect, greater than that seen with *rps28aΔ* strains. An alternative hypothesis is that the *RPS28B* mRNA, whose 3’UTR has the ability to recruit Edc3, may act as an mRNA scaffold driving PB assembly.

To assess the relative roles of Rps28 protein and *RPS28B* mRNA, we assessed PB assembly in WT, *rps28aΔ* and *rps28bΔ* strains transformed with WT *RPS28B* (*RPS28B*+3’UTR) or empty control vectors (Figure 2A). PBs form normally in the WT strain with the empty vector, with a modest increase (not significant) when a second copy of *RPS28B* is expressed from a plasmid (Figure 2A, B). In the *rps28bΔ* strain, WT *RPS28B*+3’UTR plasmid expression rescues PB assembly whereas empty vector expression does not, as expected (Figure 2A,B). Since Rps28 protein levels do not differ significantly between *rps28bΔ* strains transformed with the two constructs (Figure 2D), this argues that Rps28 protein levels are not responsible for PB assembly phenotypes in these strain backgrounds.

**Figure 2.**
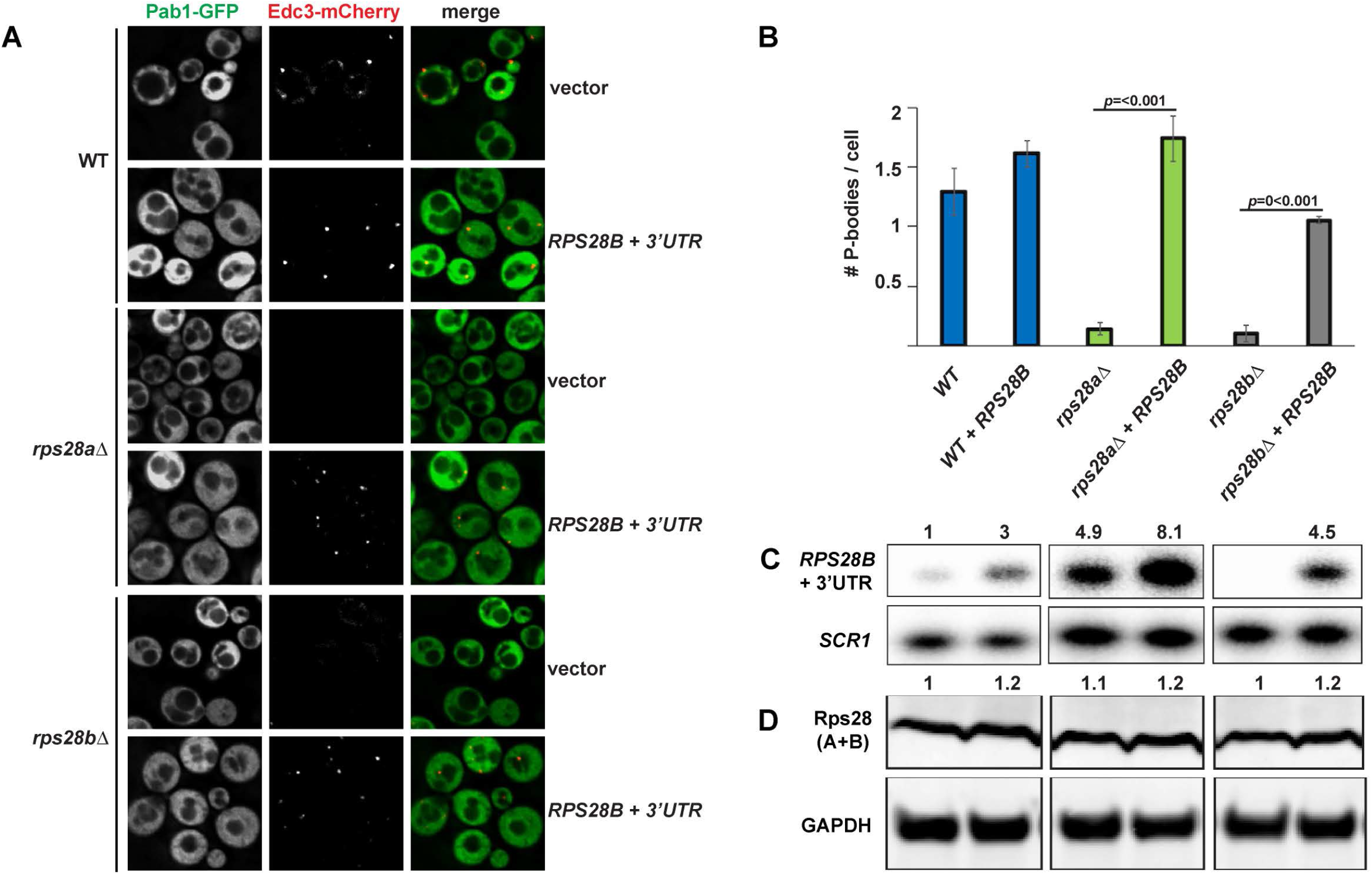
RPS28B 3’UTR is important for PB assembly. A) WT*, rps28aΔ* and *rps28bΔ* strains were transformed with empty vector, pRS315 or pRS315 expressing *RPS28B* + 3′UTR and assessed for the presence of PB foci. B) Quantitation of data in panel A. Data generated from 3 biological replicates with mean ± s.d shown. An ANOVA with Tukey’s post-hoc test was used to asses significance. C) Northern analysis of above strains with probe for RPS28B 3’UTR sequence. D) Western analysis of above strains to assess total Rps28 protein levels.

Notably, in the *rps28aΔ* strain, which already possess an endogenous *RPS28B* gene, only expression of a second copy of *RPS28B* from a plasmid rescues PB assembly while empty vector expression does not. Again, Rps28 protein levels show no discernible difference in either of the transformed *rps28aΔ* strains, whereas *RPS28B* mRNA levels are clearly increased in both *rps28aΔ* transformed strains relative to WT cells, but more so in those expressing the *RPS28B* plasmid (compare Figure 2C, lanes 3-4 versus 1). Thus, while *RPS28B* mRNA levels do alter in correlation with PB formation, suggesting a role for the *RPS28B* mRNA, there is not a correlation between overall abundance and PB assembly (i.e. no constant threshold mRNA level to induce PB formation).

One explanation to reconcile PB phenotypes with these observations of *RPS28B* mRNA and Rps28 protein abundance, and prior data in the field, is that increased abundance and/or translation of *RPS28B* mRNAs in a *rps28aΔ* strain may be compensating for the absence of *RPS28A* mRNA (compare 2C lanes 1 v 3-4). This ultimately results in normal Rps28 protein levels in both cases (Figure 2D). Importantly, given there are no obvious changes in Rps28 protein stability in WT, *rps28aΔ* or *rps28bΔ* strains (Figure S2), the differences in *RPS28B* mRNA abundance in *rps28aΔ* cells expressing either an empty vector or second *RPS28B* gene copy logically suggests that *RPS28B* mRNA must be more heavily translated in *rps28aΔ* empty vector cells. Sequestration in heavier polysomes may thus prevent effective PB scaffolding by *RPS28B* mRNA in empty vector expressing *rps28aΔ* strains, whereas non-translating *RPS28B* mRNA is more likely to exist in *rps28aΔ* cells with higher *RPS28B* mRNA abundance owing to a second *RPS28B* gene copy. Thus, the critical threshold amount of *RPS28B* mRNA necessary to form PBs may not relate to abundance, but rather the amount of *RPS28B* mRNA in a non-translating state, capable of entering into and scaffolding PBs (see discussion).

In summary, our results suggest that total Rps28 protein levels do not account for PB assembly defects in the *rps28aΔ* and *rps28bΔ* strains and suggests that *RPS28B* mRNA might be important for PB assembly.

### Truncations of the RPS28B 3’UTR can negatively and positively affect PB assembly

To more directly assess the importance of the *RPS28B* 3’UTR in PB assembly, we created 3 truncations of *RPS28B* 3’UTR and assessed their effects on PB assembly in a *rps28bΔ* strain background (Figure 3). The distal region of *RPS28B* 3’UTR (Region 3, Δ316-529) harboring the stem-loop previously reported to interact with Edc3, significantly impaired PB assembly. Deletion of the ORF-proximal region (Region 1, Δ22-179) modestly impaired PBs also, thought this effect was not significant. Nonetheless, it is formally possible that Edc3 or other PB-stimulating proteins may bind the *RPS28B* 3’UTR at more than one site (see discussion). Interestingly, deletion of region 2 (Δ180-315) significantly stimulates PB assembly. Importantly, the PB assembly defect in region 3 and 1 deletion mutants is not a result of lower *RPS28B* mRNA and protein levels; in fact both are slightly higher in these mutants than WT cells (Figure 3D,E). In summary, PB assembly can be negatively and positively modulated by truncations of the *RPS28B* 3’UTR, bolstering the argument that the *RPS28B* 3’UTR is important for PB assembly.

**Figure 3.**
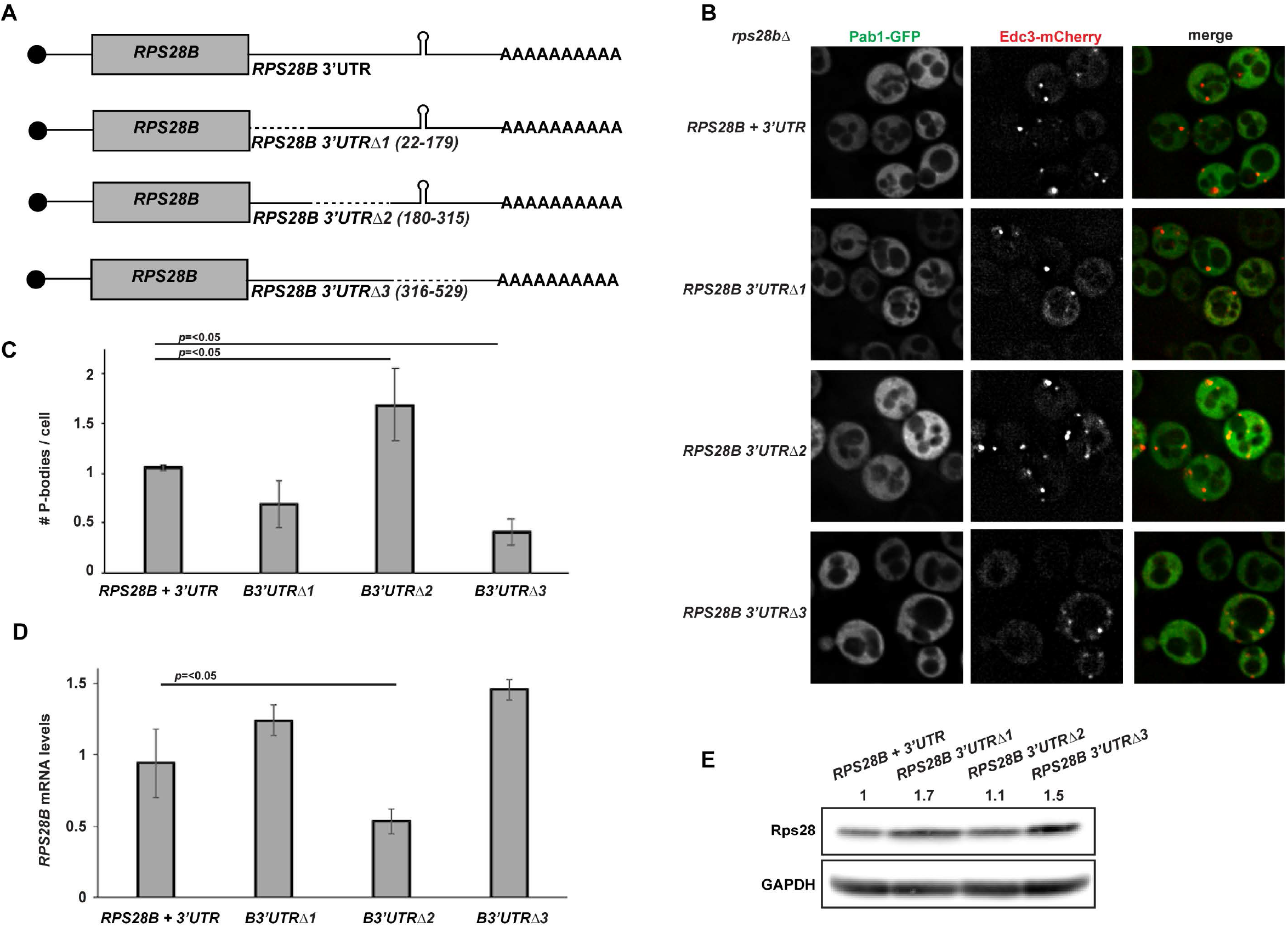
The proximal and distal RPS28B 3’UTR are important for PB formation. A) Schematic of *RPS28B* 3’UTR truncations generated. B) Log-phase *rps28bΔ* strains were transformed with plasmids harboring different *RPS28B* 3’UTR truncations and pRB001 expressing both Pab1-GFP (SG marker) and Edc3-mCh (PB marker) and assessed for their effects on PB assembly by fluorescence microscopy. C) Quantitation of data in panel B. Data generated from 3 biological replicates with mean ± s.d shown. An ANOVA with Dunnetts post-hoc test was used to assess significance. D) RT-qPCR analysis to assess *RPS28B* mRNA levels of above strains. An ANOVA with Dunnetts post-hoc test was used to assess significance. E) Western analysis of above strains to assess total Rps28 protein levels.

### RPS28B mRNA localizes to PBs

If *RPS28B* mRNA directly facilitates PB assembly, perhaps by scaffolding interactions of key PB assembly proteins, one would expect *RPS28B* mRNA to localize to PBs. To assess this, we carried out single molecule FISH. ~51% of PBs have *RPS28B* mRNA perfectly co-localized, ~25% of PBs have *RPS28B* mRNAs adjacent to them while another 24% do not have visible *RPS28B* mRNA co-localized (Figure 4). The high percent of PBs with *RPS28B* mRNA associated with them is consistent with the idea that *RPS28B* mRNA might aid PB assembly by acting as an mRNA scaffold. The 25% PBs without visible *RPS28B* mRNA might be explained by limitations of single molecule FISH (e.g. limited probe access to dense mRNP granule structures[42]; loss of mRNA from PBs during hybridization process), *RPS28B* mRNAs being in the process of degradation or that *RPS28B* mRNA is only transiently required to localize within PBs in order to stimulate PB assembly.

**Figure 4.**
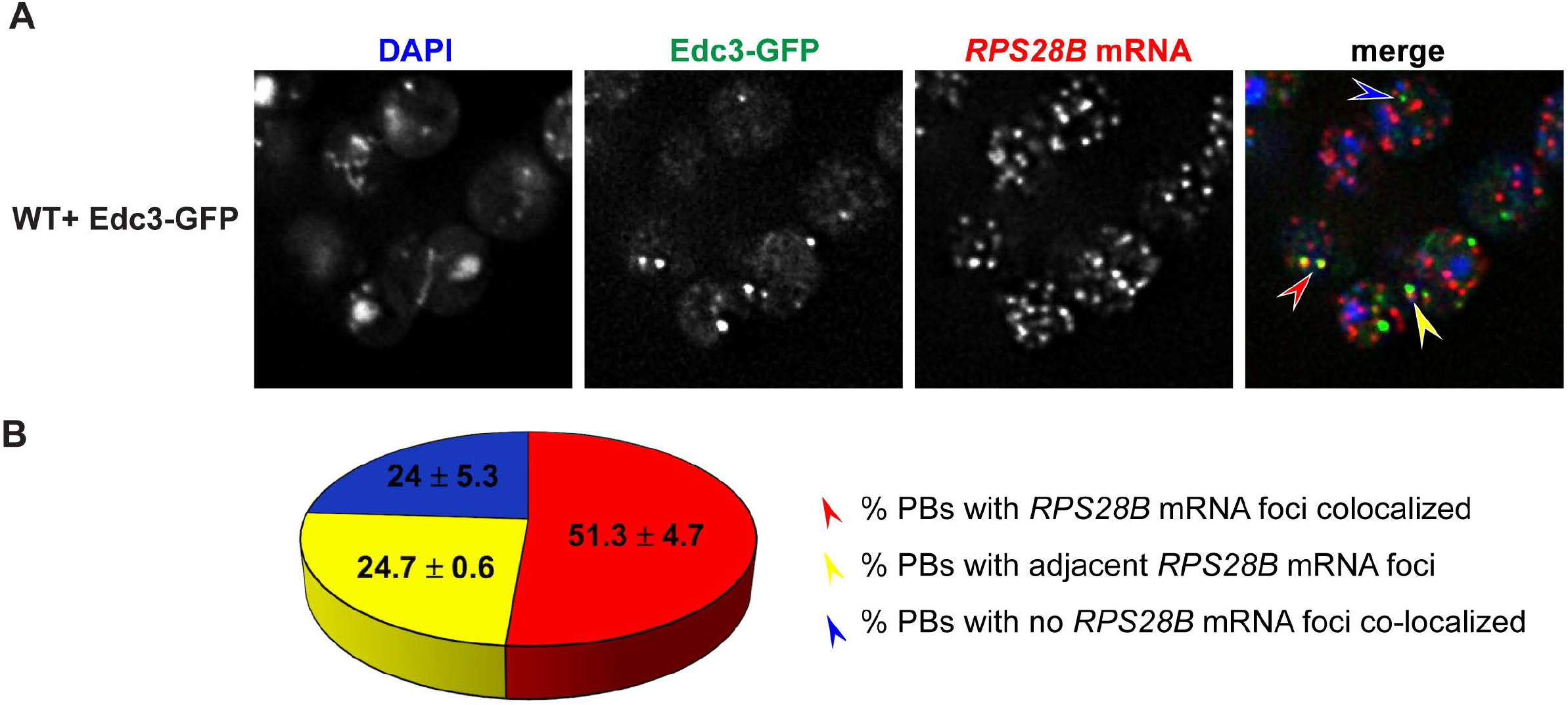
RPS28B mRNA localizes to PBs. A) WT transformed with Edc3-GFP was assessed for localization of *RPS28B* mRNA to PBs by single molecule FISH. B) Quantitation of data in A. Arrow colors in A correspond to phenotypes mentioned in B.

### Rps28 protein expressed in cis from an RPS28B 3’UTR-containing mRNA is required for PB assembly

The data presented above argues that the 3’UTR of the *RPS28B* mRNA has a significant role to play in PB assembly. However, although Rps28 protein levels did not correlate with PB assembly phenotypes (Figure 2B, D), this did not conclusively prove that *RPS28B* mRNA-based effects operated independently of the presence of Rps28 protein. Thus, to further test if the *RPS28B* 3’UTR alone could drive PB formation, chimeric constructs (Figure 5A) featuring either the *PGK1* ORF*, RPS28A* or B ORFs, or the reverse complement of the *RPS28B* ORF, fused to WT *RPS28B* 5’ and 3’UTRs, were expressed in *rps28bΔ* strains; these constructs were then assessed for their ability to rescue PB assembly.

**Figure 5.**
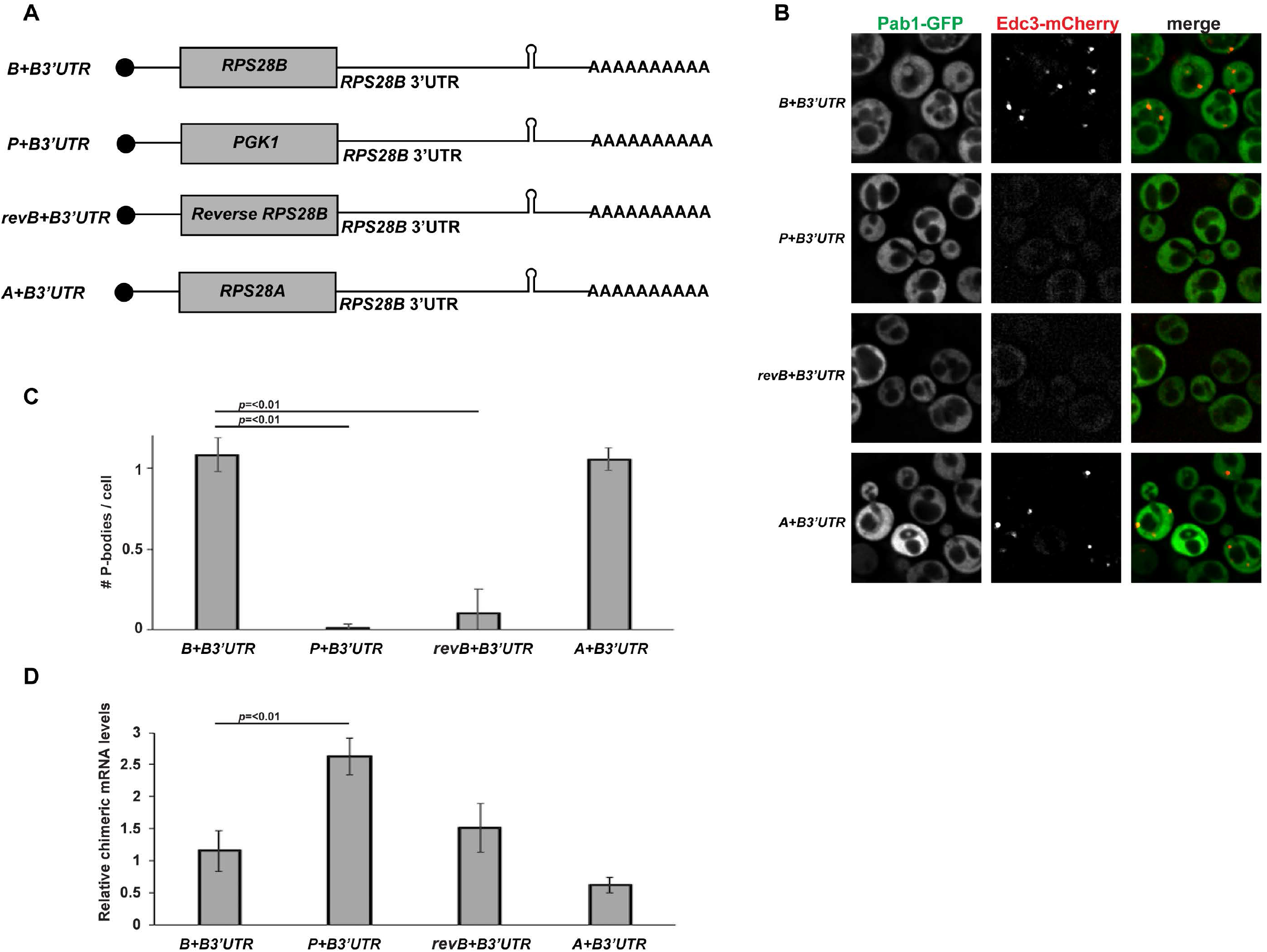
The RPS28B 3’UTR alone is not sufficient to rescue PBs in the rps28bΔ strain. A) Schematic of chimeric constructs used. B) Log-phase *rps28bΔ* strains were transformed with chimeric constructs and pRB001 and assessed for presence of PB foci. C) Data generated from 3 biological replicates with mean ± s.d shown. An ANOVA with Dunnetts post-hoc test was used to assess significance. D) RT-qPCR using *RPS28B* 3’UTR-specific probe(s) was used to assess chimeric mRNA levels. An ANOVA with Dunnetts post-hoc test was used to assess significance.

Surprisingly, we found that only the chimeric mRNA expressing *RPS28A* or *RPS28B* ORFs fused to *RPS28B* 3’UTR were sufficient to fully rescue PB assembly in *rps28bΔ* strains (Figure 5B-C). *PGK1* or reverse complemented *RPS28B* ORFs fused to the *RPS28B* 3’UTR failed to stimulate PB assembly in the *rps28bΔ* background. These results were not artefacts of altered expression levels of the chimeric mRNAs; in fact, strains lacking PBs typically expressed higher levels of their *RPS28B*+3’UTR containing chimeric mRNAs than cells expressing fully WT *RPS28B* mRNA, where PBs did form (Figure 5D). Interestingly, the *RPS28A+RPS28B* 3’UTR construct was previously shown to undergo decay similar to WT *RPS28B+RPS28B* 3’UTR while the other chimeric mRNAs fail to be degraded as efficiently[17], suggesting a possible relationship between *RPS28B* mRNA decay and PB assembly (see discussion). Additionally, expressing start codon mutants of otherwise WT *RPS28B*+3’UTR constructs in a *rps28bΔ* background failed to rescue PBs (data not shown). The simplest explanation given these results is that translation of Rps28a or b protein *in cis* from an mRNA harboring the *RPS28B* 3’UTR is critical for PB assembly.

### Rps28 protein translated in cis from an RPS28B 3’UTR containing mRNA facilitates interaction of nascent Rps28 protein with Edc3

Based on prior Edc3-Rps28 and Edc3-*RPS28B* 3’UTR interaction studies[17–19] and the above data, we hypothesized that “*cis*-translation” of Rps28 protein from an *RPS28B* 3’UTR-harboring mRNA facilitates interaction of nascent Rps28 with 3’ UTR-bound Edc3 (Figure 6A), which in turn enhances PB assembly via an unclear mechanism.

**Figure 6.**
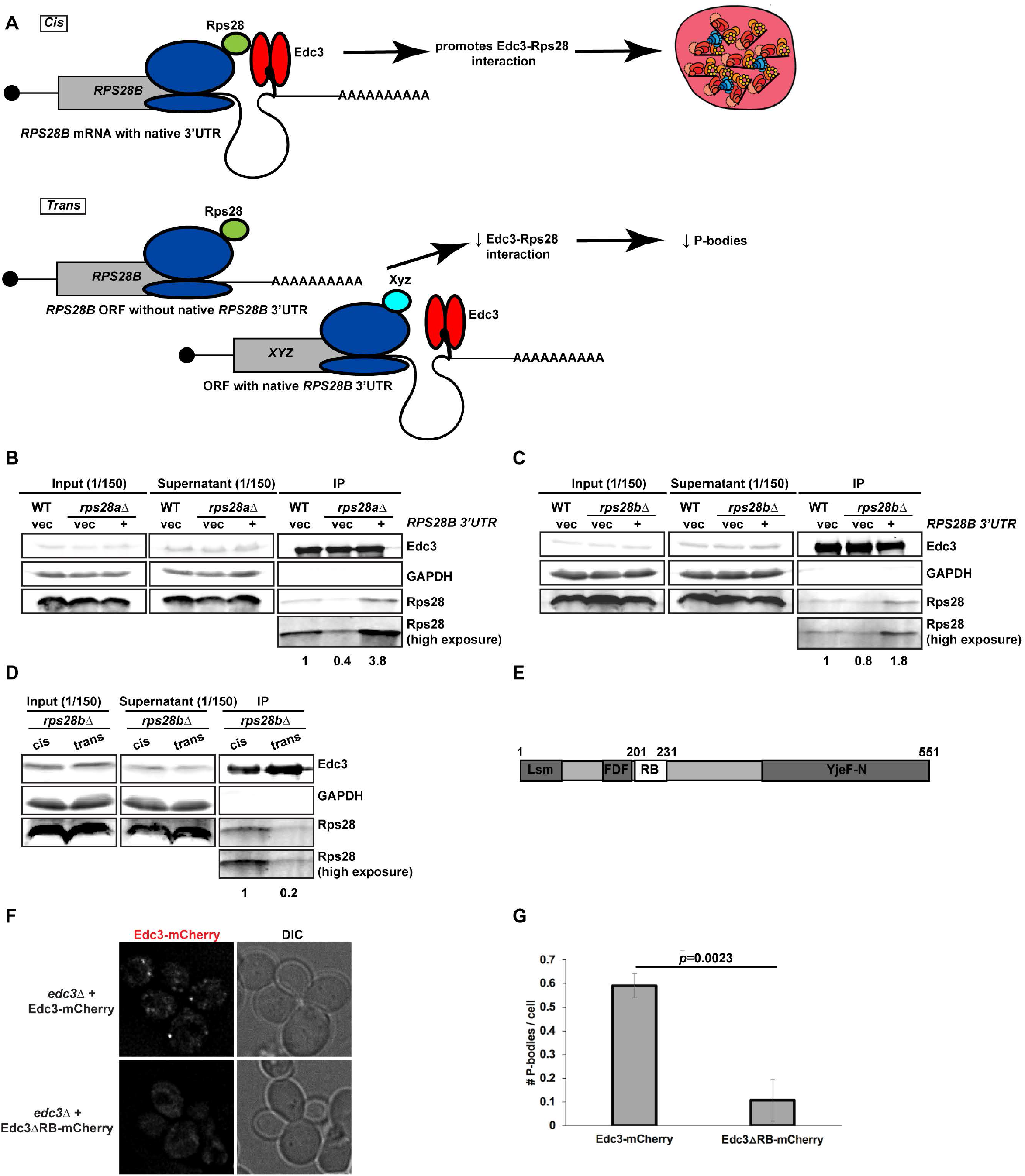
Rps28 protein translated in cis from the RPS28B mRNA facilitates an Rps28-Edc3 interaction. A) Working model of nascently synthesized Rps28 interacting better with Edc3 if Rps28 ORF lies immediately upstream of Edc3-bound *RPS28B* 3’UTR, which in turn aids PB assembly. B,C) Log-phase WT, *rps28aΔ* and *rps28bΔ* strains transformed with empty vector and *RPS28B* were assessed for Rps28-Edc3 interaction by carrying out an IP for Edc3-GFP and probing for Rps28. Rps28 IP exposure is same duration as in input/supernatant lanes; longer Rps28 exposures are also shown to better quantify relative differences in Edc3-Rps28 interaction. Quantification of Rps28 band intensity is normalized to Edc3 IP band intensity. Images/quantification representative of 2 biological replicates. D) “*cis*” *rps28bΔ* yeast transformed with *RPS28B* ORF-*RPS28B* 3’UTR and “empty vector” (tTA plasmid - pRB079); “trans” *rps28bΔ* yeast transformed with *PGK1* ORF-*RPS28B* 3’UTR and *RPS28* ORF-*CYC1* 3’UTR, tTA (pRB388). Images, quantification and biological replicates as in B-C. E,F) Log-phase *edc3Δ* RP840 strains transformed with Edc3-mCherry/*Edc3ΔRB*-mCherry were assessed for Edc3-mCherry PB foci by microscopy. G.) Quantification of data in F; data was analyzed via a 2-tailed student’s T-test.

To test this hypothesis, we first determined if Edc3 can bind the *RPS28B* 3’UTR directly, as previous Yeast-3-hybrid (Y3H) data[17] did not fully rule out the possibility of a bridging protein being involved. We conducted an Electrophoretic Mobility shift assay (EMSA) using purified Edc3 and a 60mer oligo encapsulating the *RPS28B* 3’UTR stem loop region and confirmed that Edc3 can indeed bind the *RPS28B* 3’UTR directly (Figure S3); interestingly, a stepped pattern suggests multiple Edc3 binding events at higher concentrations may occur. Next, we carried out an immunoprecipitation of Edc3-GFP and assessed its ability to interact with Rps28 protein in a WT strain expressing an empty vector, and an *rps28aΔ* or *rps28bΔ* strain expressing either an empty vector or a WT *RPS28B* construct (Figure 6B,C), which our prior data indicated rescued PBs (Figure 2). Interestingly, Rps28 protein interaction with Edc3 was barely detectable in *rps28aΔ* or *rps28bΔ* strains expressing empty vectors but rose significantly in WT cells and *rps28aΔ* or *rps28bΔ* strains expressing WT *RPS28B* constructs. Thus, increased Rps28-Edc3 interaction correlates with PB assembly. This suggests that the *RPS28B* 3’UTR facilitates interaction of nascently-translated Rps28 protein with Edc3. A simple model is that this *in cis* 3’UTR-driven protein-protein interaction reflects the closer spatial proximity of Rps28 synthesis to 3’UTR-bound Edc3 than in the scenario of an *RPS28* ORF being separated from its endogenous 3’UTR (Figure 6A). To directly test this, we examined the effect of supplying *rps28bΔ* cells with a transcript bearing the *RPS28B* ORF and *RPS28B 3’UTR* in *cis*, or where the *RPS28B* ORF and 3’UTR were supplied *in trans* on separate transcripts (Figure 6D). Strikingly, despite equal Edc3 and Rps28 protein levels, we again saw a stronger interaction of Edc3 and Rps28 protein in cells bearing the *cis RPS28B* ORF and 3’UTR transcript. Similar results were observed in *rps28bΔ* strains using the WT and *PGK1* chimeric plasmids (B+B3’UTR and P+B3’UTR) outlined in Figure 5A (Figure S4). Collectively, this data strongly argues that the presence of *RPS28B* ORF and 3’UTR in an *rps28bΔ* strain only facilitates a robust Edc3-Rps28 protein interaction when both these elements are on the same transcript.

Interestingly, if proximity of Edc3 to nascently synthesized Rps28 is indeed key to their interaction, this might also explain the increase in PBs with truncation 2 that deletes a region in the middle of the *RPS28B* 3’UTR (Figure 3B,C). Deletion of this region would result in bringing the *RPS28B* 3’UTR region 3 with the stem-loop (Edc3 binding site) closer to the *RPS28B* ORF, possibly facilitating the Rps28-Edc3 interaction.

In summary, in both *rps28bΔ* and *rps28aΔ* backgrounds, the degree to which PBs form correlates with the ability of Edc3 and Rps28 protein to immunoprecipitated, which occurs most strongly when the *RPS28B* ORF and 3’UTR are in *cis*.

### The Rps28-Edc3 protein interaction facilitates PB assembly

To determine if the Rps28-Edc3 protein interaction indeed facilitates PB assembly, we turned to a previously described mutant of Edc3 impaired in this interaction[19]. Using *edc3Δ* cells expressing plasmid-borne WT Edc3-mCherry or the Rps28 binding mutant (Edc3ΔRB-mCherry; Figure 6E)[19], we observed that PB assembly is strongly inhibited in Edc3ΔRB-mCherry expressing cells compared to cells expressing WT Edc3-mCherry (Figure 6F-G). Thus, the Edc3-Rps28 protein interaction is indeed important for PB assembly.

## Discussion

Previous work has emphasized protein-protein interactions as key drivers of PB and RNA granule assembly in general[10,43,44][45]. However, given that RNAs can base pair with other RNA molecules and bind to numerous RNA-binding proteins, they too are clearly multivalent and thus, perhaps unsurprisingly, also play a key role in RNA granule assembly. Supporting this, treating cells with cycloheximide completely inhibits PB and SG assembly due to sequestration of mRNA in polysomes, whereas puromycin treatment stimulates PBs and SGs due to release of mRNAs from polysomes [9,46,47]. Additionally, biochemical evidence indicates that RNAs can drive liquid-liquid phase separation (LLPS) of granule proteins *in vitro* [24]. *In vivo*, RNA promotes phase separation of Meg proteins in P-granules *Caenorhabditis elegans* [48]. Finally, yeast total RNA can self-assemble *in vitro*, in the absence of any proteins, into phase separated granules whose RNA content closely resembles the transcriptome of *in vivo* purified SGs; this suggests that RNA-RNA interactions are often likely to be sufficient to recruit RNAs to SGs and drive their assembly[24]. While RNA is clearly a driver of RNP granule assembly, specific RNAs (with the exception of the lncRNA NEAT1 in paraspeckles[49]) have not been identified as scaffolding the assembly of RNP granules. This study is the first to our knowledge showing that a specific yeast mRNA, *RPS28B,* drives PB assembly.

Several features of *RPS28B* mRNA make it a good candidate for scaffolding PBs. First, the *RPS28B* mRNA 3’UTR is one of the longest 3’UTRs in yeast, about 643 nucleotides long with the median yeast 3’UTR being ~120 nucleotides long[50]. Second, in WT cells, *RPS28B* mRNA is seemingly translated weakly compared to its paralog *RPS28A*, making it more available to scaffold PBs which only harbor non-translating mRNAs[51]. Supporting this, previous data and our data (not shown) suggests that while *RPS28A* mRNA is only 50% more abundant than *RPS28B* at steady state in WT cells[18], Rps28a protein is ~11 fold higher than Rps28b protein in WT cells[20], indicating a large difference in translational efficiency. Furthermore, ribosome profiling data indicates that *RPS28A* is translated more efficiently than *RPS28B*[51]. Third, the *RPS28B* 3’UTR directly binds (Figure S3; [17]) to Edc3, one of the major yeast PB assembly proteins. Given that our *RPS28B* truncation data (Figure 3A-C) hints that more than 1 site in the 3’UTR may contribute to PB assembly, it is also quite possible that the *RPS28B* 3’UTR might be a binding site for other protein interactions that drive PB assembly. Indeed CLIP data previously showed that Dhh1 binds to *RPS28B* mRNA but not *RPS28A* mRNA[52]. Dhh1 is a core PB component previously implicated in repression of translation initiation, elongation and stimulating mRNA decapping [35,53]. Alternatively, Edc3 may also bind at more than 1 *RPSB28B* 3’UTR site. Regardless, the ability of *RPS28B* mRNA to drive PB assembly in yeast raises an intriguing question; do other specific RNAs, mRNA or otherwise, drive the formation of other RNA granules? In addition to genetic and phenotypic screening, recent advances in purifying RNA granules, coupled with sequencing of their RNA content, may help reveal the answer to this question.

*RPS28B* mRNA is a well-studied example of autoregulatory control of ribosomal protein production which our study adds new insight to and raises new questions. Work by the Jaquier, Seraphin and Jacobson labs suggests that when Rps28 protein (A or B) levels are in excess, an Edc3-dependent but deadenylation-independent rapid decay of *RPS28B* mRNA occurs. A stem loop structure in the *RPS28B* 3’UTR appears critical to this Edc3-facilitated mRNA turnover[18]. The importance of the Edc3-Rps28 binding interaction to *RPS28B* mRNA turnover is less clear, with the Seraphin lab describing a necessity for this interaction for Edc3-facilitated *RPS28B* mRNA decay[19], whereas no effect was seen by the Jacobson lab on steady state *RPS28B* mRNA levels[17]. In our study, we find that overall Rps28 protein levels are remarkably well controlled, with essentially no variation in overall abundance in WT, *rps28aΔ* or *rps28bΔ* strains that do or do not express a WT copy of *RPS28B* on a plasmid (Figure 2), nor an alteration in Rps28 protein stability in WT, *rps28aΔ* or *rps28bΔ* strains (Figure S2). However, *RPS28B* mRNA levels do vary significantly, with *RPS28B* steady state levels strongly increasing in *rps28aΔ* nulls (Figure 2) as expected. Determining whether *RPS28A* mRNA is subject to autoregulation is an area of future interest, especially given that this transcript seemingly accounts for the majority of total Rps28 protein in WT cells. It is also intriguing that *rps28bΔ,* but not *rps28aΔ* strains exhibit a modest growth defect (Figure S1F), suggesting a possible functional importance for *RPS28B* beyond simply production of Rps28 protein.

The ability to form *RPS28B*-stimulated PBs seems to rely on similar elements reported as being required for forming an Edc3-decay competent *RPS28B* mRNP. An *RPS28* ORF needs to be upstream of the *RPS28B* 3’UTR in a *rps28bΔ* background ([17]; and Figure 1), and an interaction between Rps28 and Edc3 is required ([19]; and Figure 6). Intuitively, it may seem odd that an mRNA subject to accelerated decay could also facilitate assembly of PBs. However, it is possible that the *RPS28B* mRNA merely plays a transitory role in stimulating the assembly of a Rps28-Edc3 protein complex, and that this complex, and PBs themselves may persist after the *RPS28B* mRNA itself is degraded. This would be consistent with our smFISH data (Figure 4).

A simple model for the *RPS28B* 3’UTR scaffolding PB assembly predicts that it’s presence alone in cells would be sufficient to stimulate RNA and/or protein interactions that drive PB formation; to our surprise, this was not the case. PB assembly requires an *RPS28* ORF (A or B) upstream of the *RPS28B* 3’UTR (Figure 5A-B). Additionally, a previously described Rps28-Edc3 protein interaction[17,19] is strengthened when the *RPS28B* ORF is upstream of the *RPS28B* 3’UTR (Figure 6B-D). Finally, impairing Rps28-Edc3 protein interaction directly with an Edc3 Rps28 binding mutant also impairs PB assembly (Figure 6E-G). Note, this differs from findings by the Seraphin lab[19], who reported no effects on PB assembly in cells expressing WT or Rps28 binding-mutant Edc3; however quantitative data was not presented, and a second copy of Dcp2 was also expressed in their system which may conceivably alter PB assembly thresholds given that Dcp2 interacts with Edc3 and other PB proteins. Regardless, the above observations suggest that translation of Rps28 protein directly upstream of the *RPS28B* 3’UTR increases the probability of binding Edc3 (a *cis*-translational interaction), and that this interaction in turn stimulated PB assembly. How Rps28-Edc3 protein interaction achieves this is unclear; indeed, different evidence has been presented for existence of heterodimeric and trimeric (2 Edc3:1 Rps28) protein complexes *in vitro* and *in vivo* respectively[17,19]. This remains an area of future study but nonetheless represents another example of a ribosomal protein with an intriguing extra-ribosomal function. The fact that this function happens to be facilitating the assembly of PBs, whose assembly is anti-correlated with bulk translation levels[5,35], suggests a possible control point for balancing general translation activity with translation repression and decay in yeast.

A role for 3’UTRs driving formation of functionally important protein interactions, in which the *cis*-translated protein is one of the protein interacting partners has recently been described for two mRNA 3’UTRs in human cells by the Mayr lab[26–28]. In one example, an extended 3’UTR isoform of the *CD47* gene (a cell-surface ‘marker of self’ protein), specifically recruits the RNA binding protein HuR which in turn recruits SET. CD47 translated in *cis* in turn interacts with SET, which ultimately enhances CD47 localization to the plasma membrane [26]. In contrast, a short 3’UTR isoform of the CD47 transcript fails to recruit HuR and SET, causing CD47 to preferentially localize to the ER, where CD47 translation occurs. In the other example, mass spectrometry analyses revealed that the *BIRC3* gene, an E3 Ubiquitin ligase, when encoded from a long 3’UTR isoform (but not a short 3’UTR isoform), leads to formation of many distinct BIRC3-containing protein complexes[28]. These included BIRC3 forming a complex with protein trafficking factors IQGAP and RALA, which are themselves recruited to the *BIRC3* long 3’UTR by RNA binding proteins Staufen and HuR. Ultimately the *BIRC3* long 3’UTR, and the resulting BIRC3-IQGAP-RALA protein complex facilitates recycling to the cell surface of receptor proteins CXCR4 and CD27. In a *BIRC3* long 3’UTR isoform null context, impaired CXCR4 membrane localization likely underpins an observed cell migration defect in a malignant B cell model system [28]. Thus, the Mayr lab studies and our own illustrate two key principles. First, 3’UTRs may not just serve as regulators of mRNA function, but also as facilitators of protein-protein interactions with broad-reaching functional consequences for the protein interactors. Secondly, the effects of 3’UTRs on promoting consequential protein-protein interactions seem to depend on *cis*-translation for one of the protein interactors, and recruitment of the other protein interactor(s) to the 3’UTR. Our work demonstrates this phenomenon extends from yeast to mammals and can happen in very different biological contexts (cell surface protein localization versus PB assembly), and as a result of differing gene expression regulatory mechanisms (alternatively spliced 3’UTRs versus differentially expressed gene paralogs). A key open question is how widespread the role of 3’UTRs and *cis*-translation is in facilitating functional protein-protein interactions.

## Acknowledgements

We are grateful to the Jacquier, Badis, Jacobson and Séraphin Lab for plasmids, and the Parker Lab for plasmids and purified Edc3 protein; we particularly acknowledge Bhalchandra Rao and Laura Mizoue for their help here. This work was supported by startup funds to J.R.B. from the University of Arizona and support from NIGMS (NIH RO1-GM1145664).

---

**Supplemental Figure 1.**
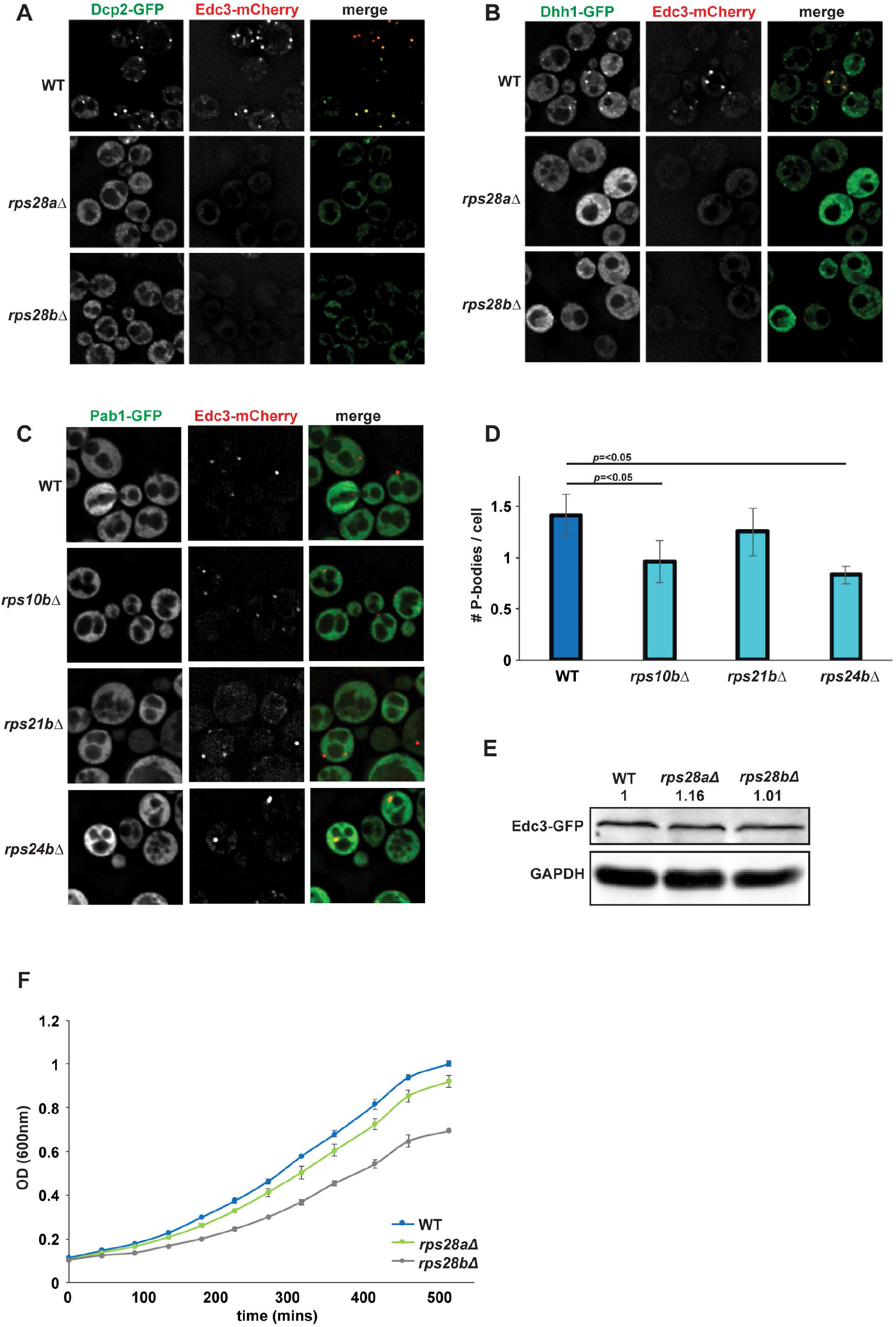
RPS28A and RPS28B are important for PB assembly. A,B) The PB defect in the *rps28aΔ* and *rps28bΔ* strains is consistent when analyzed by other PB markers. Logphase WT, *rps28aΔ and rps28bΔ* strains transformed with plasmids expressing Dcp2-GFP (pRB038) or Dhh1-GFP (pRB038) and Edc3-mCherry (PB markers) were examined for the presence of PB foci. C,D) Other ribosomal protein deletion mutants do not have a severe defect in PB assembly. Log-phase WT*, rps10aΔ, rps10bΔ, rps21aΔ, rps21bΔ* and *rps24bΔ* strains were assessed for their ability to form PBs as in A. Average number of PBs per cell. Data generated from 3 biological replicates with mean ± s.d shown. Paired 2-tailed student T-tests were used to assess significance. E) Levels of the PB marker Edc3-GFP do not differ significantly in WT, *rps28aΔ* and *rps28bΔ* strains. Log-phase WT, *rps28aΔ* and *rps28bΔ* strains transformed with Edc3-GFP (pRB224) were assessed for Edc3-GFP levels by western blotting. F) A growth curve was carried out to assess growth of WT, *rps28aΔ and rps28bΔ* strains. Data generated from 3 biological replicates with mean ± s.d shown.

**Supplemental Figure 2.**
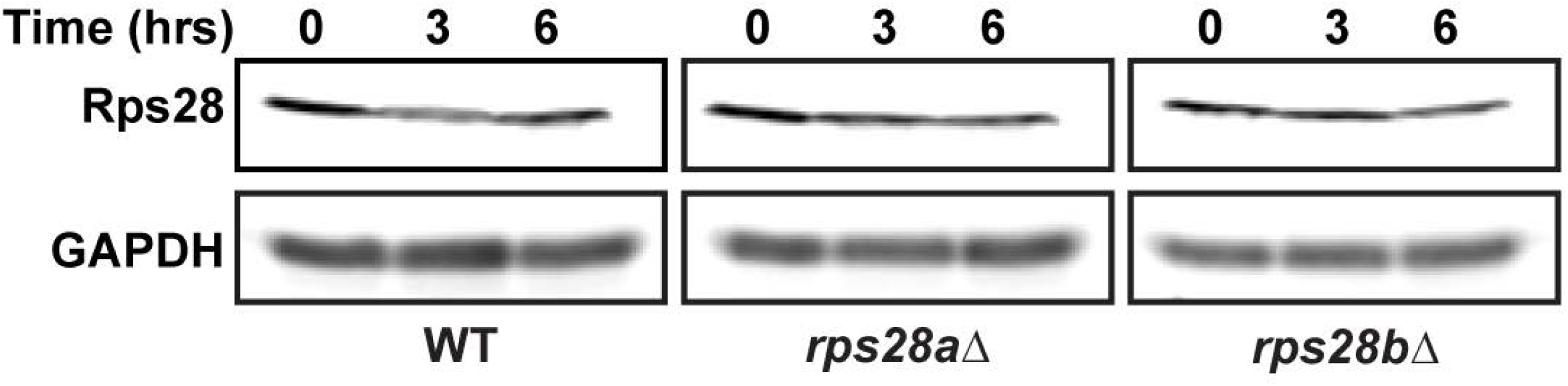
Rps28 protein stability does not vary in WT, rps28aΔ or rps28bΔ backgrounds. Turnover of Rps28 protein in the indicated strains was carried out by using a cycloheximide (100μg/ml) shut-off experiment.

**Supplemental Figure 3.**
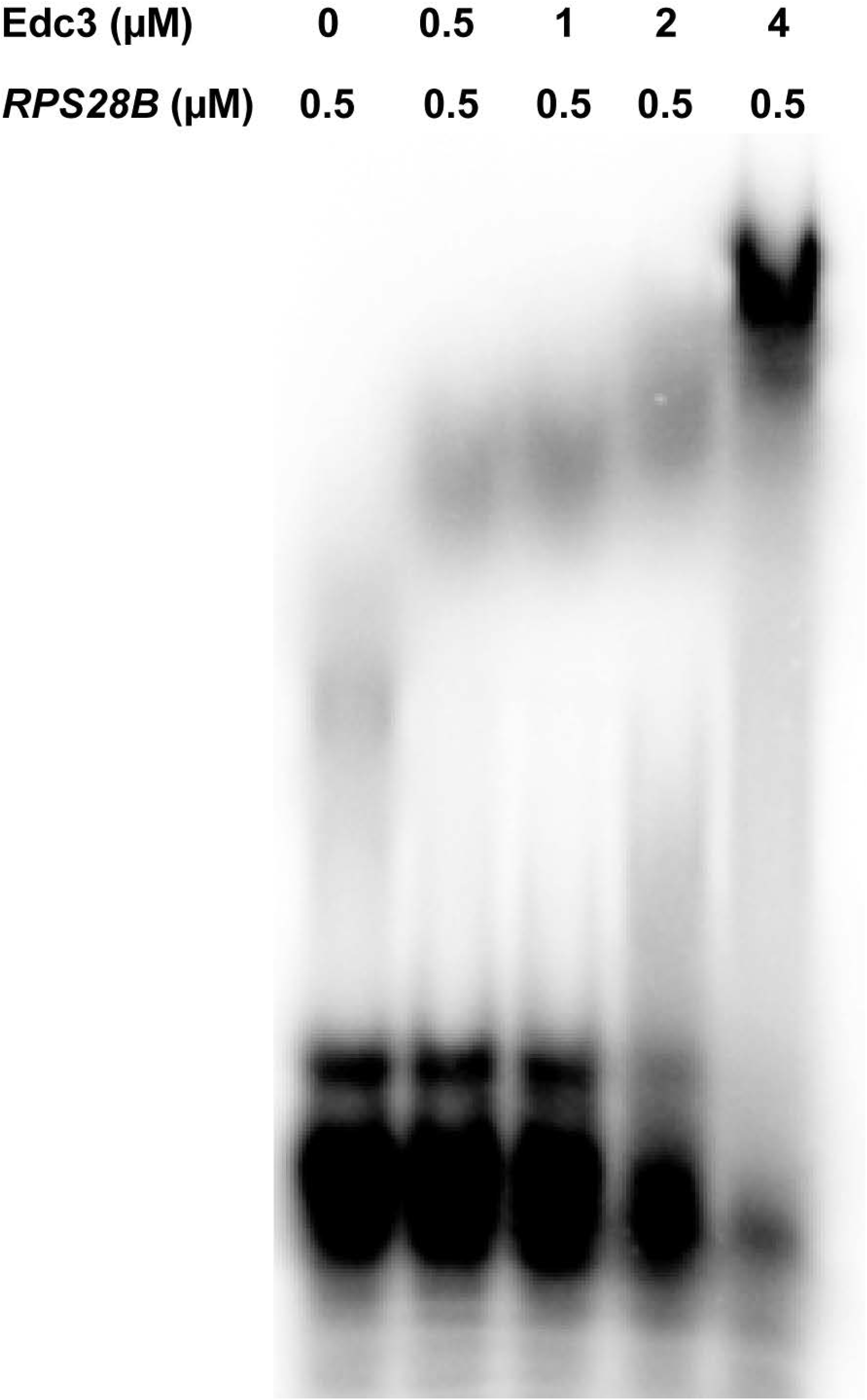
Edc3 directly interacts with the stem-loop structure in RPS28B 3’UTR. EMSA was carried out using Edc3 protein and a synthesized 60nt *RPS28B* RNA oligo containing the predicted *RPS28B* stem-loop structure. The indicated concentration of Edc3 was mixed with ^32^P end-labeled RNA (0.5μM). Following incubation at 30°C for 60 min, the reaction mixtures were resolved on 6% native PAGE gel.

**Supplemental Figure 4.**
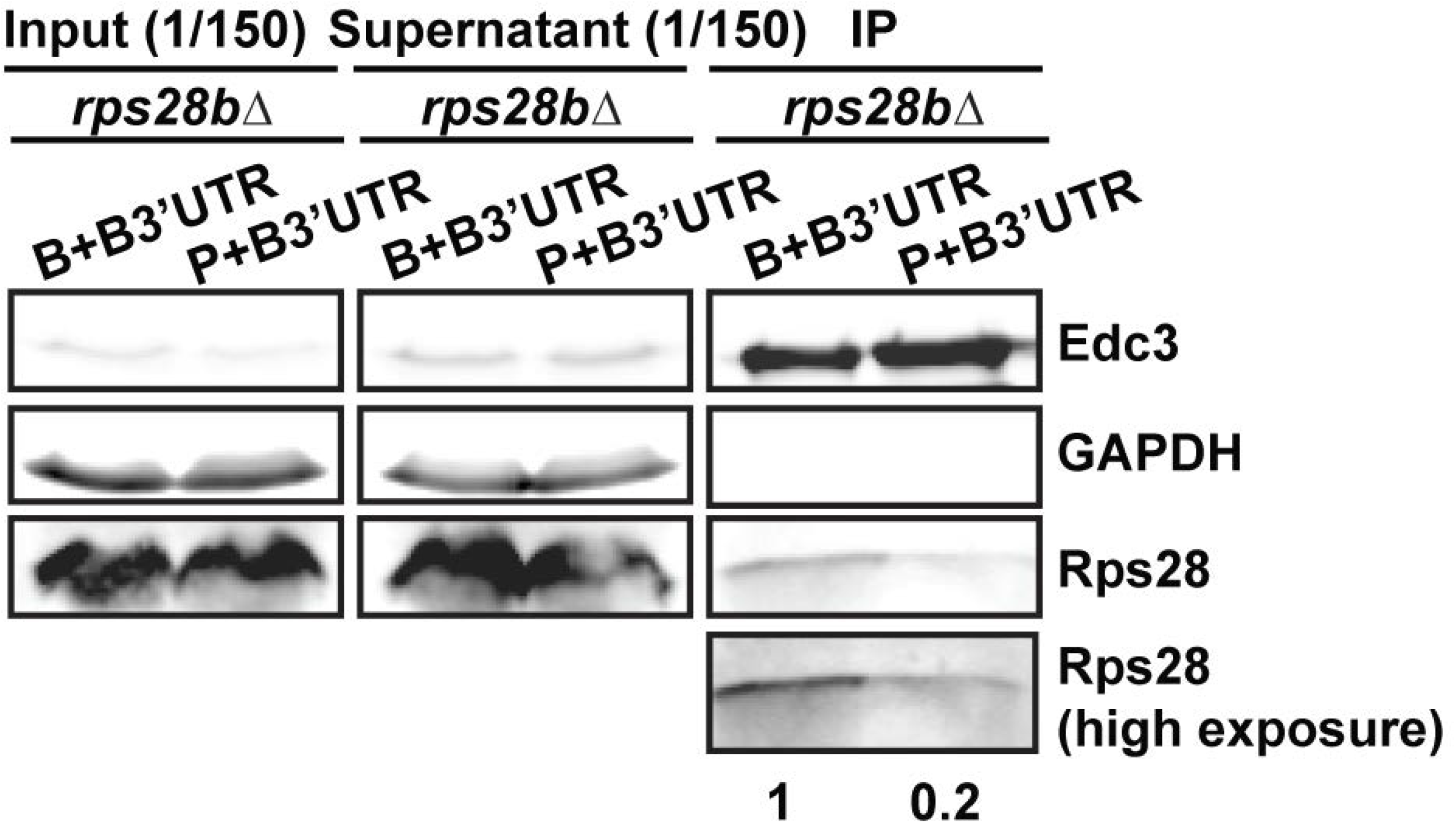
Rps28 protein translated in cis from the RPS28B mRNA facilitates an Rps28-Edc3 interaction. Log-phase *rps28bΔ* strains transformed with *RPS28B* ORF-*RPS28B* 3’UTR and *PGK1* ORF-*RPS28B* 3’UTR chimera and assessed for Rps28-Edc3 interaction as in Figure 6B and C

**Supplementary Data 1:**
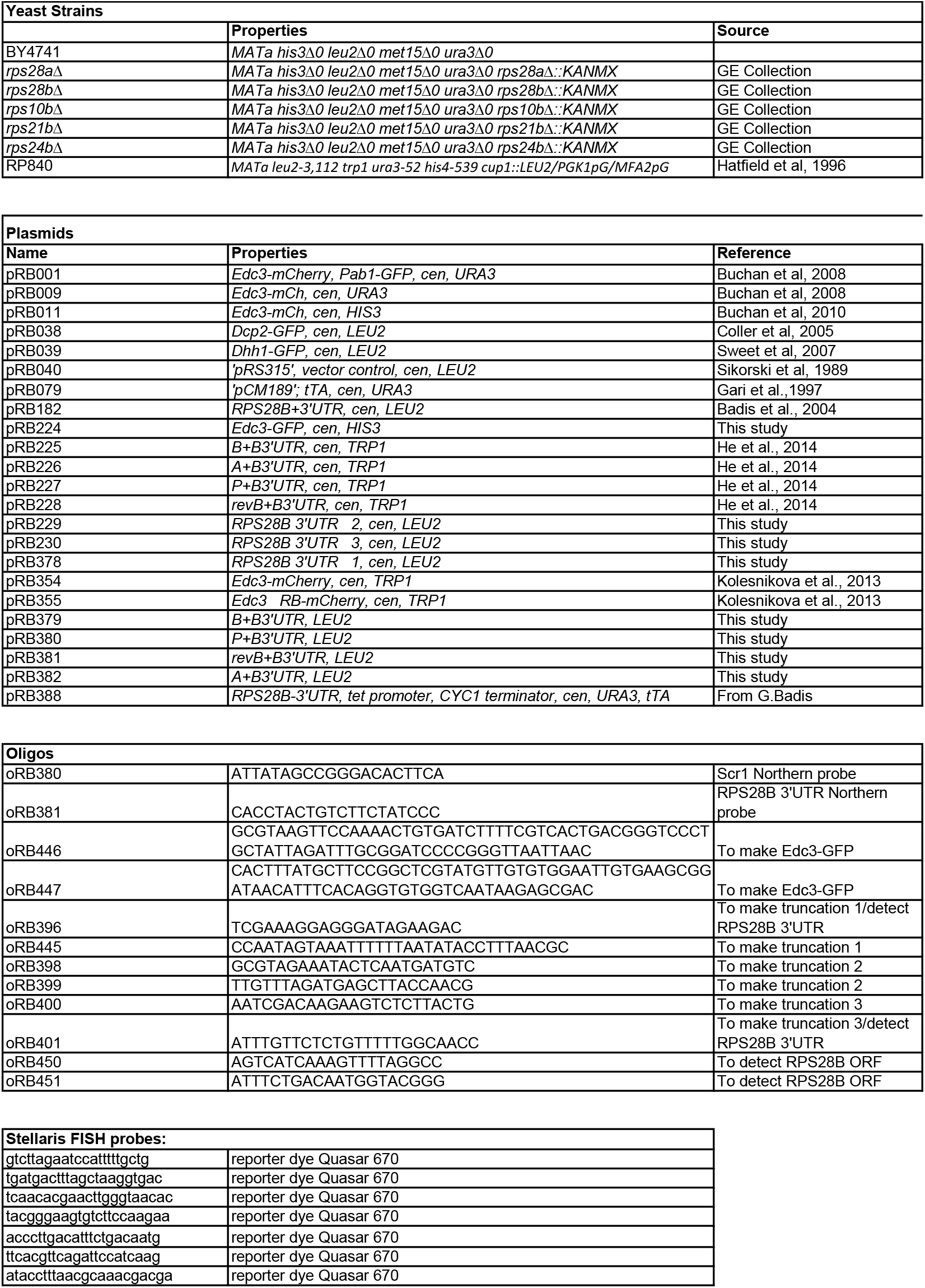
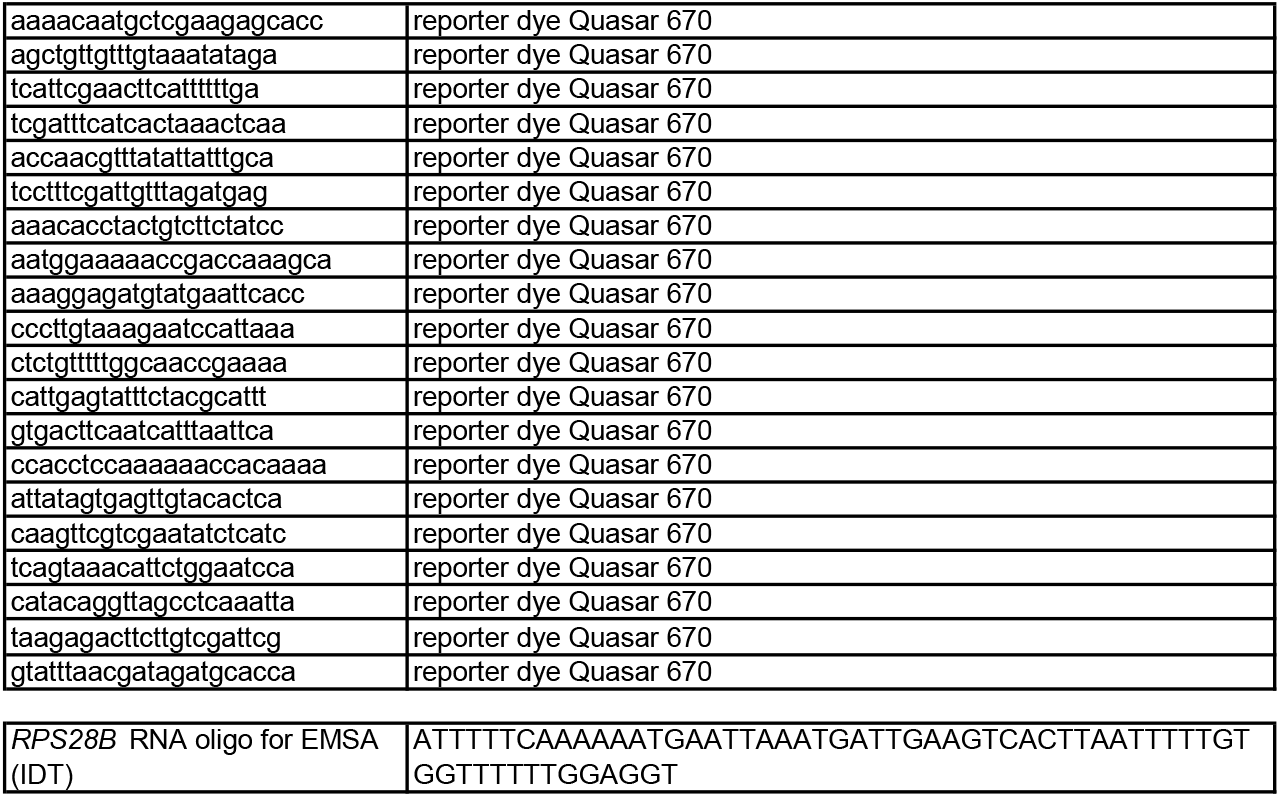
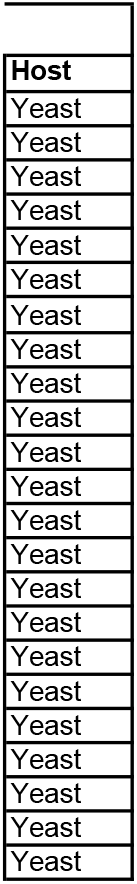
Strains, Plasmids and oligos used in this study.

